# Voxel-wise regression without intensity normalization provides better sensitivity when identifying injury regions after stroke in FDG-PET images

**DOI:** 10.1101/2023.08.23.554408

**Authors:** Wuxian He, Hongtu Tang, Jia Li, Xiaoyan Shen, Xuechen Zhang, Chenrui Li, Huafeng Liu, Weichuan Yu

## Abstract

**Background:** In the analysis of brain fluorodeoxyglucose positron emission tomography (FDG-PET), intensity normalization is a necessary step to reduce inter-subject variability. However, the choice of an appropriate normalization method in stroke studies remains unclear, leading to inconsistent findings in the literature.

**Materials and methods:** Here, we propose a regression-based and single-subject-based model for analyzing FDG-PET images without intensity normalization. Two independent data sets were collected before and after middle cerebral artery occlusion (MCAO), with one comprising 120 rats and the other comprising 96 rats. After data preprocessing, voxel intensities in the same region and hemisphere were paired before and after the MCAO scan. A linear regression model was applied to the paired data, and the coefficient of determination was calculated to measure the linearity. The results between the ipsilateral and contralateral hemispheres were compared, and significant regions were defined as those having reduced linearity. Our method was compared with existing intensity normalization methods and validated using the triphenyl tetrazolium chloride (TTC) staining data.

**Results:** The proposed method detected more injury areas compared to existing approaches, as confirmed by the ground truth provided by TTC. The area under the curve (AUC) of the average receiver operating characteristic (ROC) curves using our method reached 0.84, whereas the AUCs using existing methods ranged from 0.77 ∼ 0.79. The average false positive rate (FPR) and true positive rate (TPR) of the individual analysis results using our method (FPR = 0.06, TPR = 0.56) were better than the group-wise analysis results using *t*-tests (FPR = 0.10, TPR = 0.51). The identified injury regions were consistent in the two independent data sets. Some of them were confirmed by other publications.

**Conclusions:** The proposed method offers a new quantitative approach to analyzing FDG-PET images. The calculation does not involve intensity normalization and can be applied to a single subject. The method yields more sensitive results than existing intensity normalization methods.

## 1. Introduction

[^18^F]fluorodeoxyglucose (^18^F-FDG) positron emission tomography (PET) is a widely used nuclear medicine imaging technique to study glucose consumption in animals and human patients (Fu et al., 2009; Zimmer & Luxen, 2012). It is particularly useful in the study of neurological disorders, such as Alzheimer’s disease (AD) (Chételat et al., 2020; Mosconi et al., 2009), Parkinson’s disease (Eckert et al., 2005; Matthews et al., 2018), dementia (Kato et al., 2016; Shivamurthy et al., 2015), and stroke (Bunevicius et al., 2013; Lee et al., 2021), where altered glucose metabolism can be identified in the brain. However, the absolute value of the image intensities in FDG-PET imaging is not an accurate representation of the glucose uptake level due to disease-unrelated and patient-dependent factors, such as FDG concentration, plasma glucose level, and basal metabolic rate (Boellaard, 2009; Claeys et al., 2010; Dukart et al., 2010). Therefore, image intensity normalization becomes a necessary step for further quantitative evaluation of the FDG-PET images.

Different intensity normalization methods have been proposed and used, with the most prevalent one being proportional scaling using the cerebral global mean (CGM) (Frackowiak et al., 2004). This method assumes that only a small area in the brain has changed and that the between-group differences in the CGM are negligible. It has been adopted in the popular analysis program Statistical Parametric Mapping (Penny et al., 2011) and other software programs (Jeong et al., 2019) to discriminate mild neurological disorders from healthy controls. However, for brain diseases with large dysfunctional regions, such as AD and stroke, intensity normalization using CGM may produce extensive false positives (Yakushev et al., 2009). To exclude the altered regions and improve the robustness, a single reference region believed to have minimal influence can be used (Küntzelmann et al., 2013). Regions such as the cerebellum and pons are commonly used as reference regions in studies of diseases with known mechanisms. However, in stroke studies where the location of ischemia and infarction is not entirely recognized, a well-accepted reference region is still lacking (Nie et al., 2018). To address this issue, more data-driven methods have been proposed. For example, Andersson (Andersson, 1997) and Nie et al. (Nie et al., 2018) proposed iterative approaches by repeatedly performing proportional scaling and excluding altered areas. Yakushev et al. (Yakushev et al., 2009) utilized the “apparent hypermetabolism” obtained from global mean normalization which reached higher sensitivity. Histogram-based normalization is another type of data-driven method that normalized PET images in a voxel-to-voxel manner to an averaged template so that voxel intensities could be compared directly between different scans (Proesmans et al., 2021).

Despite the diversity of intensity normalization strategies, there is currently no consensus on how to choose a proper intensity normalization method to study a certain disease. For instance, in stroke studies, various normalization approaches have been used, including proportional scaling by the global mean (Nie et al., 2014), the contralateral hemisphere (Yuan et al., 2013), and iterative methods (Nie et al., 2018). However, different intensity normalization methods yield quite different identification results, and even falsely detected voxels from distinct methods also have large variations (Lopez-Gonzalez et al., 2020; Nie et al., 2018). Consequently, there is substantial inconsistency in the findings regarding the location of significant regions. For instance, some researchers only observed hypo-metabolism (reduced glucose uptake) in the ipsilateral hemisphere (Li et al., 2020; Sobrado et al., 2011), whereas other researchers revealed a hypo-uptake pattern in the ischemic core as well as hyper-uptake (increased glucose uptake) regions in the penumbra (Arnberg et al., 2015; Yuan et al., 2013). It is difficult to determine why such inconsistency occurs because both normalization strategies and batch effects can bring variability. Therefore, a robust and reliable method to analyze FDG-PET images is desirable.

An additional constraint of current approaches is their reliance on group-level comparisons, such as the two-sample *t*-test. This necessitates the establishment of a detection threshold (typically *α* = 0.05 following adjustment) and the collection of an adequate number of samples (Borghammer et al., 2009) to achieve statistical significance. As a result, the finding of the analysis is restricted to a group-level framework and the overall outcome can only be explained in the context of group comparisons. Consequently, it is not possible to assess the results at an individual level. Nevertheless, a method that can directly analyze individual samples would be considerably advantageous.

In this paper, we present a novel regression-based approach to analyzing FDG-PET images without intensity normalization. The method can be applied to a single subject and can identify regions of injury following brain ischemia. The approach quantifies the linearity of voxel intensity values in the same region before and after an ischemic stroke. Specifically, brain regions that exhibit low correlation with the disease are expected to have better coherence in the paired intensities. In contrast, brain regions with poor linearity of paired intensities before and after the stroke are more likely to be dysfunctional following the ischemia. To assess the efficacy of the proposed method, data from two independent rat middle cerebral artery occlusion (MCAO) experiments were used. The performance of the algorithm was evaluated using 2,3,5-triphenyl tetrazolium chloride (TTC) staining and compared with existing approaches using receiver operating characteristic (ROC) analysis. The results of the comparison suggest that the proposed approach has the highest sensitivity among all benchmark methods for identifying injury regions following stroke.

## 2. Materials and methods

### 2.1. Subjects

All the animal subjects in this study were adult female Sprague-Dawley rats weighing 180–280 g. The experiment consisted of two groups of rats, one comprising 120 rats and the other including 96 rats. The rats underwent right brain MCAO surgery using the same methodology as outlined in (Longa et al., 1989), but at different time points, resulting in two independent data sets. Throughout the experiments, all subjects were provided with free access to water and food, except for an overnight fast before PET scanning. On the day of the surgery, MCAO was performed on the right brain of each rat to induce cerebral ischemia. During the MCAO procedure, the animal was placed on a heating pad to maintain its temperature. The skin was incised using a scissor, and the temporalis muscles were retracted, exposing the zygomatic and squamosal bones. A high-speed drill was used to create a small hole (2-3 mm) at the fusion of the zygomatic and squamosal bones. The middle cerebral artery on the right brain was identified and exposed by retracting the dura using an ophthalmic scissor and forceps. After that, the artery was permanently ligated by bipolar electrical coagulation. Prior to the MCAO surgery and PET scan, chloral hydrate (400 mg/kg) was administered as an anesthetic. The anesthetized animals were monitored throughout the experiment, and the chloral hydrate was supplemented as needed. This study was approved by the Animal Research Committee of the School of Medicine, Zhejiang University. All the procedures complied with the ARRIVE guidelines and were carried out in accordance with the National Institutes of Health guide for the care and use of Laboratory animals (NIH Publications No. 8023, revised 1978).

### 2.2. FDG PET

All PET images were acquired in 3D mode using a microPET R4 scanner (CTI Concorde Microsystems, LLC.) located at the Medical PET Center of Zhejiang University. The scanner has a spatial resolution of 1.88 mm full width at the half maximum (FWHM) in the axial plane and 1.9 mm FWHM in the transverse plane. Each rat received an injection of FDG with a concentration of 0.5-1 mCi from the tail vein, followed by a 15-minute PET scan performed 30 minutes later. After five hours, the radiolabeled glucose in the rat’s body was mostly attenuated and consumed. The MCAO surgery was then performed, and another dose of 0.5-1 mCi FDG was injected 15 min after the procedure. The second acquisition was performed under the same conditions as the baseline scan, collecting metabolic information after the ischemic stroke.

### 2.3. Image preprocessing and analysis

The raw data were reconstructed as 128 × 128 × 63 volumes of 1 × 1 × 1 mm^3^ voxel size using the vendor’s software, which adopted the maximum a posteriori algorithm with 18 iterations (*β* = 0.5 resolution) (G. Wang et al., 2008). After reconstruction, spatial preprocessing of the acquired images was performed using statistical parametric mapping software (SPM12, Wellcome Department of Imaging Neuroscience, London, United Kingdom) embedded in MATLAB R2018b (MathWorks Inc., Natick, MA). The images were subjected to scaling, alignment, segmentation, and spatial normalization using an in-house FDG PET template for Sprague-Dawley rats, following the same methodology as outlined in our previously published work (He et al., 2022). The normalized images only contained the intracranial portion of the brain and were smoothed with a Gaussian kernel of 2 × 2 × 4 mm^3^ FWHM. The final data were presented in an 80 × 108 × 63 volume of 2 × 2 × 2 mm^3^ voxel size.

All data analyses were performed using MATLAB R2018b. A volumetric rat brain atlas in the Waxholm space was initially utilized for region delineation (Papp et al., 2014). The atlas was comprised of 222 regions of interest (ROI) represented by the voxel coordinates. It was spatially normalized to the FDG PET template, thereby registering the atlas with the PET images.

### 2.4. Motivation of the method

The proposed method was motivated by an observation of the correlation between the intensity levels of baseline and MCAO scans of the same subject. The striatum, specifically the caudate putamen, was examined as an example since it is a well-established ischemic core following a stroke. Fig. 1 illustrates this relationship in three rats, where the left column represents the contralateral left brain. The voxel intensities display a high degree of linearity between the baseline and MCAO scan, which is confirmed by the coefficient of determination *R*^2^. In contrast, the right column exhibits voxel intensities that are poorly linear and randomly distributed. The corresponding *R*^2^ indicates a low level of predictability of MCAO intensities based on the baseline intensities. The MCAO procedure and the resultant alteration of FDG concentration in the ipsilateral brain are responsible for the significant difference in linear relationships between the two hemispheres.

**Figure 1:**
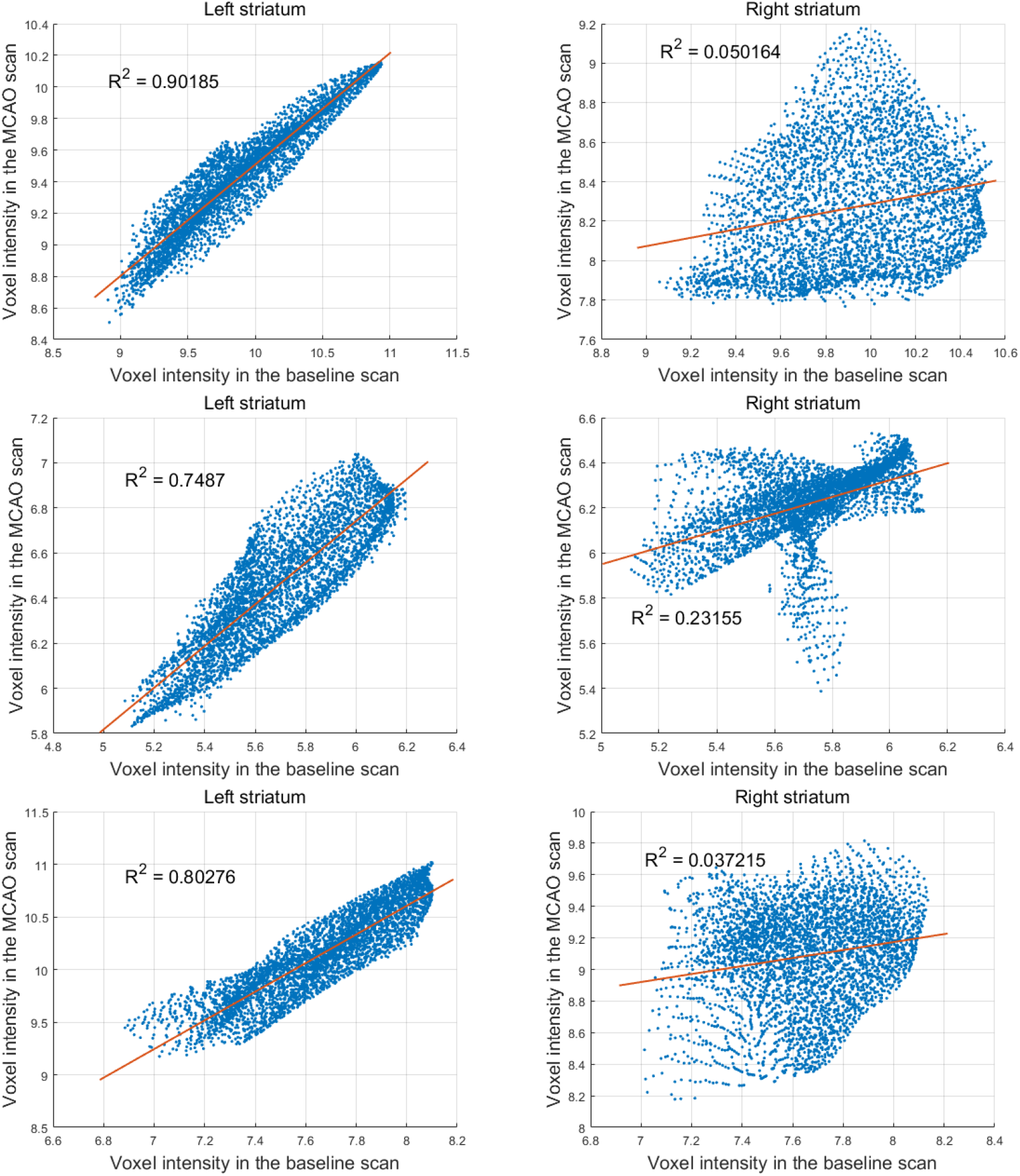
The scatter plots of voxel intensities in the striatum between the baseline and MCAO images of three representative rats, with each row containing data for one rat. Least square regression was employed to fit a straight line, and the coefficient of determination *R*^2^ was computed, indicating the intensity coherence between the two images. The left column shows the results in the left hemisphere, while the right column presents data for the right hemisphere.

### 2.5. Proposed algorithm

Since each animal underwent two scans, one before and one after MCAO, and both scans were registered, the paired images were used to construct the regression model. The volumetric atlas was initially applied to each image to generate 222 sets of voxels representing 222 ROIs in both hemispheres. Linear regression models were then applied to the voxel intensities of each ROI from the image pairs.

Consider an ROI with voxel intensity variables (*x*_*i*_, *y*_*i*_), *i* = 1, …, *n* where *x*_*i*_’s represent the voxel intensities in the first (baseline) scan, *y*_*i*_’s represent the voxel intensities in the second (MCAO) scan, and *n* is the number of voxels within the ROI. The regression model fits a straight line using the least-square algorithm by minimizing the sum of squared residuals *S* between the two variables:

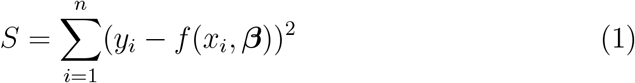

where *f* (*x*_*i*_, ***β***) is the fitted line with *f* (*x*_*i*_, ***β***) = *β*_0_ + *β*_1_*x*_*i*_. Two parameters need to be estimated from the data with *β*_0_ being the *y*-intercept and *β*_1_ being the slope.

After the line was fitted, the coefficient of determination *R*^2^ of the regression model was computed. *R*^2^ is defined as the proportion of variation in the dependent variable (*y*_*i*_’s) that can be predicted by the independent variable (*x*_*i*_’s) and can be calculated through the formula:

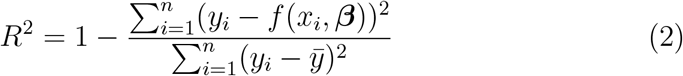

where 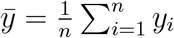. The value of *R*^2^ ranges from 0 to 1, with higher values indicating greater predictability of the model.

Upon obtaining the *R*^2^ values, the subsequent step involved analyzing the results of each ROI in both hemispheres. If an ROI displayed a large *R*^2^ value close to 1, and this pattern was consistently observed in most samples, it would be categorized as a *stable* region. Conversely, if one region exhibited a significantly smaller *R*^2^ value in the right hemisphere compared to the left hemisphere, and this pattern was consistently observed among most subjects, it was expected to have been disturbed or damaged by the MCAO. Consequently, this ROI would be categorized as an *unstable* region. The difference in *R*^2^ values between the left and right hemispheres (left - right) was computed for all ROIs and rats. Subsequently, the ROIs with a median greater than a threshold were chosen as the final unstable regions.

### 2.6. Benchmarking with existing intensity normalization

The proposed regression model was compared with existing intensity normalization methods. In this study, we selected and implemented the following four approaches:

- Global mean normalization (GMN): It divided each voxel by the mean intensity over the whole brain. The voxels inside the brain parenchyma were segmented with a brain mask, which was produced from the template or averaged images (He et al., 2022).
- Reference region normalization (RRN): It divided each voxel by the mean intensity over a certain region believed to have minimal influence. In this study, three commonly used reference regions from the contralateral hemisphere were evaluated including the brain stem (Lopez-Gonzalez et al., 2020; Nugent et al., 2020), the cerebellum (Dukart et al., 2010; Küntzelmann et al., 2013), and the pontine nuclei (Marcoux et al., 2018; Verger et al., 2021). The hypo- and hyper-uptake volumes detected by each reference region were evaluated using the Dice similarity coefficient (*DSC*) (T. Zhang et al., 2022) between the two data sets

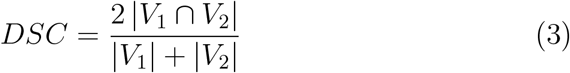

where *V*_1_ and *V*_2_ are the volumes to be compared. The reference region with the highest *DSC* was adopted for final comparison to the proposed method.
- Histogram-based normalization (HN): It divided each image to the averaged template in a voxel-to-voxel manner, producing a ratio image (Proesmans et al., 2021). Then the histogram of the ratio image was generated and the value of the highest bin was adopted as the normalizing factor. By doing so, every image was normalized to the intensity range of the template and thus can be compared horizontally.
- Cluster-based normalization (CN): It was a data-driven normalization method proposed by Yakushev et al. (Yakushev et al., 2009). First, the algorithm looked for hyper-uptake areas using GMN. Second, the hyper-uptake area with the highest *t*-value was recognized as the cluster. Finally, the mean intensity over the cluster was utilized as the new normalization factor to extract the significant regions.

### 2.7. Performance evaluation

The performance of the proposed algorithm, along with the existing methods was evaluated based on the TTC staining results (Popp et al., 2009). Eleven rats were sacrificed after the entire experiment. Their brain sections were stained with TTC to reveal the infarction, and their corresponding PET images were assessed by the proposed algorithm and compared with the existing methods. Specifically, the 2D TTC staining photos were first digitized and segmented to provide the ground truth of the injury (see Appendix A for more details). Secondly, the detected significant volume of the same rat from the 3D PET images was overlaid onto the template, and the corresponding coronal slice was located. Thirdly, a screenshot of the coronal view was saved and adjusted to the same size as the TTC results. Fourthly, the injury area indicated by the TTC was treated as the ground truth, and the detected regions were compared to find the false positives and true positives. Finally, different thresholds were tested, and the ROC curve was generated to compare the performance of the five approaches for each of the 11 rats.

The ROC curve illustrated the ability of a binary classifier with varying threshold values by plotting its true positive rate (TPR) against the false positive rate (FPR). In our setting, the TTC staining results provided the ground truth. Therefore, a significant pixel detected by a specific model became a true positive (TP) if the TTC also showed infarction at the same location. On the other hand, a false positive (FP) pixel implied that it was significant but did not have evidence from the TTC staining results. Moreover, an insignificant pixel showing infarction on the TTC was categorized as a false negative (FN), while an insignificant and non-infarct pixel became a true negative (TN). Following these definitions, we calculated the FPR and TPR according to the standard formulas (Fawcett, 2006):

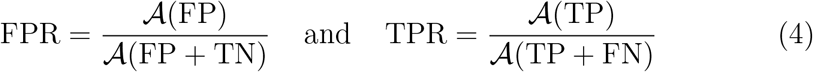

where 𝒜(·) is the area function. An averaged ROC curve was also computed using threshold averaging.

## 3. Results

### 3.1. Performance

#### 3.1.1. TTC ground truth

Following segmentation and binarization of the TTC data, the brain slice with the most prominent and largest white area was selected as the reference for infarction. Subsequently, the infarct region was extracted using a freehand ROI in MATLAB. Concurrently, unstable regions were computed from the corresponding PET images for each of the 11 subjects. A threshold of 0.2 was employed as it yielded outcomes that were most comparable to the ground truth with respect to the injury area. The 3D volume of the unstable region was superimposed onto the PET template space, and the coronal view at the same anterior/posterior (AP) coordinates as the TTC slice was saved for validation purposes. Fig. 2 shows the original and segmented TTC results (with infarction shown in white), in addition to the location of the unstable regions in the corresponding PET images. Each pair of images was aligned so that the brain slices were registered. These results will be utilized for further quantitative analysis, including comparison with existing intensity normalization techniques.

**Figure 2:**
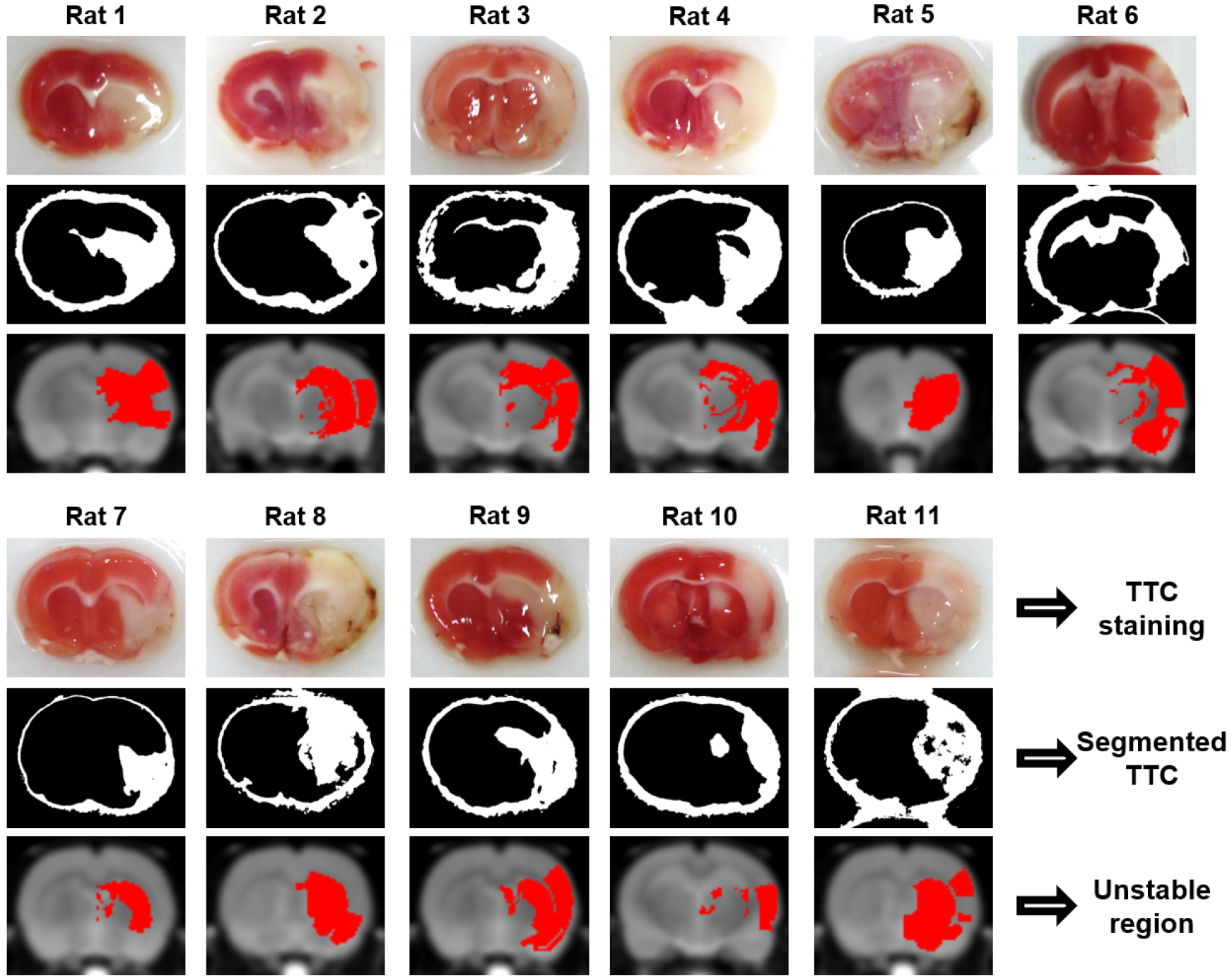
The 11 rats from the first data set with TTC staining results (first row) and the corresponding unstable regions (third row). The TTC slice was segmented from the original image, and the white area (excluding the brain boundary) was identified as the infarction (second row). The unstable region slice had the same AP coordinate as the corresponding TTC, and the red area represented its location.

#### 3.1.2. Optimal reference region

In order to compare different approaches, it is necessary to establish a fixed reference region for the RRN method. As there is no consensus on the ideal reference region for stroke studies, we selected three typical reference regions (Lopez-Gonzalez et al., 2020) and evaluated their performances. A suitable reference region should exhibit both fine intra- and inter-set consistency. To this end, we randomly divided each dataset into three equal-sized subsets and assessed the consistency of the significant volumes detected from them. Specifically, we employed each candidate as the reference region for intensity normalization and conducted the voxel-wise analysis. The two-sample *t*-test was applied with a significance level of *p* = 0.05 (family-wise error corrected), and hypo-uptake clusters with sizes larger than 20 voxels were identified (Ocklenburg et al., 2016). Hypothesis testing was carried out for each subset, resulting in three significant volumes for each data set. The DSC was then computed between every two subsets to measure consistency. The results are presented in the bar plots in Fig. 3. Both the intra- and inter-set consistency demonstrate that the cerebellum is the optimal reference region for this study.

**Figure 3:**
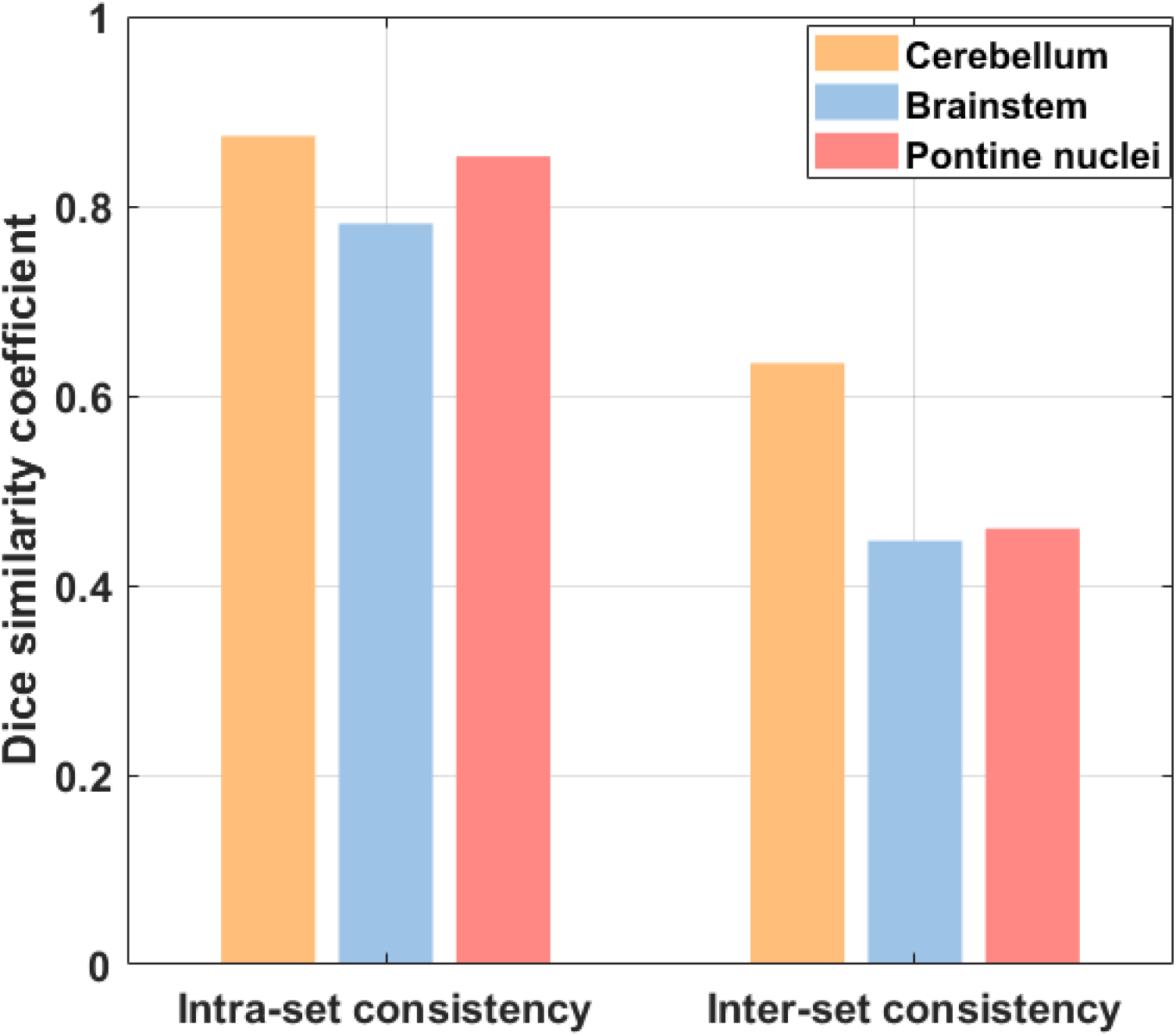
The three reference region candidates were evaluated by performing intensity normalization using each of them and checking the hypo-uptake volumes. The hypo-uptake volumes were computed for each subgroup from the two data sets. The dice similarity coefficient was adopted to measure the consistency of the results. The mean of the six intra-set consistency 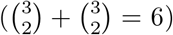 and nine inter-set consistency (3 *×* 3 = 9) were calculated. The candidate with the highest consistency was chosen to be the final reference region.

#### 3.1.3. FDG-PET quantification in existing methods

Once the cerebellum was established as the optimal choice for RRN, we proceeded to compare it with other approaches, including the proposed regression method. Each method was applied to the entire data set to determine the *t* threshold for hypo-uptake patterns (with a significance level of *p* = 0.05 and family-wise error corrected). The hypo-metabolic volume was then calculated for each of the 11 rats with TTC ground truth based on the *t* threshold. The hypo-metabolic region resulting from each normalization strategy, along with the unstable region, was compared based on the TTC infarction (Lou et al., 2007). Fig. 4. (a) shows the location of significant regions derived from the five methods for one exemplary rat. As discussed in the previous work by Borghammer et al. (Borghammer et al., 2009), a more conservative threshold of *p* = 0.001 was adopted for the CN method to prevent nearly all right brains from showing hypo-uptake patterns.

**Figure 4:**
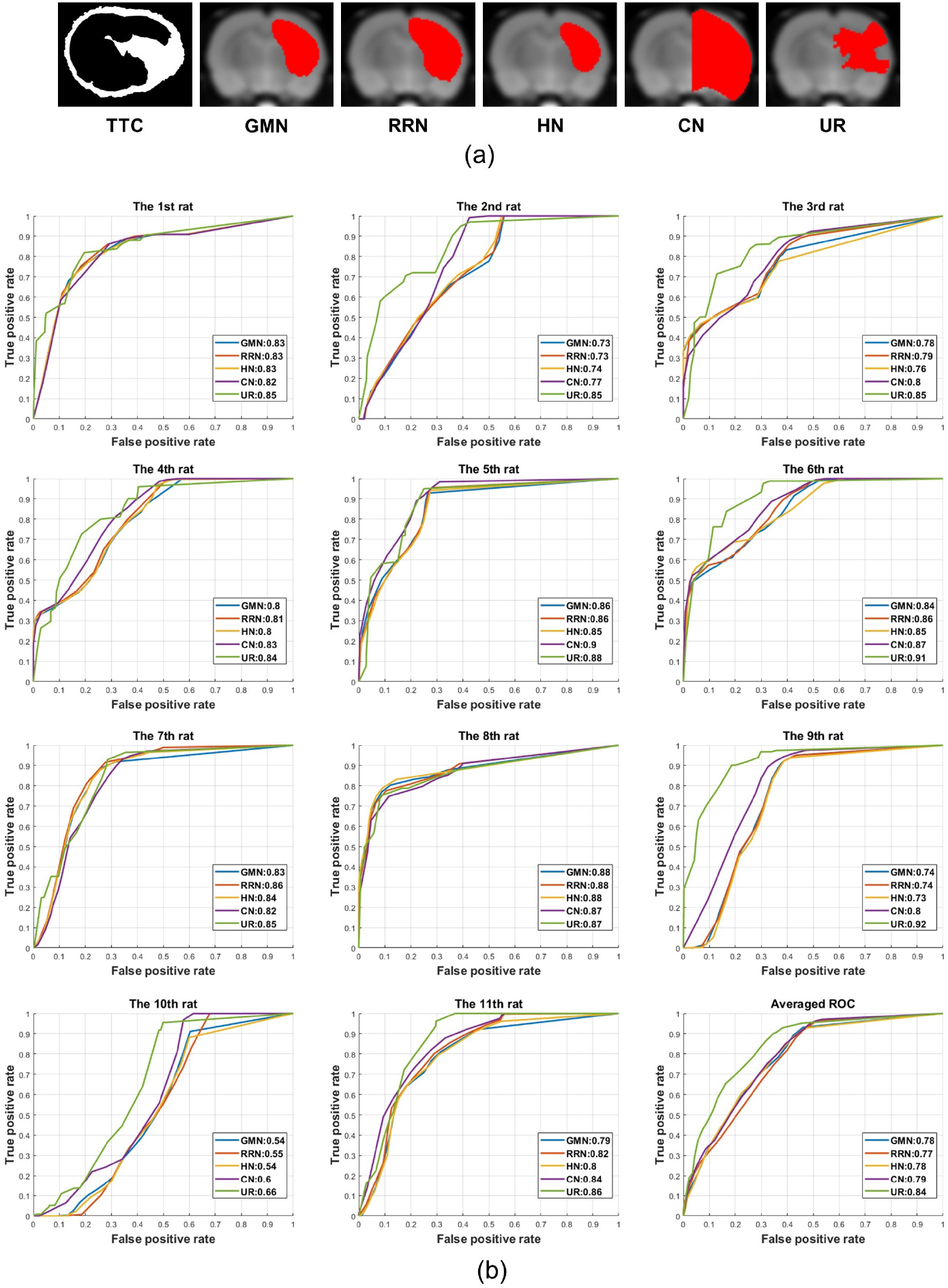
The performance of different intensity normalization methods was evaluated using ROC curves. Each method was implemented on all 11 rats with TTC ground truth, and the hypo-uptake volume computed from the PET images was superimposed onto the template. An example of the results is presented in (a). Then the ROC curve was constructed, and the AUC was calculated and displayed in the figure legends. The ROC of all 11 individuals was averaged to create an overall ROC curve using threshold averaging. GMN: global mean normalization, RRN: reference mean normalization, HN: histogram-based normalization, CN: cluster-based normalization, UR: unstable region.

#### 3.1.4. Comparison results

The ROC curves and their corresponding area under the curve (AUC) for the 11 rats are displayed separately in Fig. 4. (b). The final graph shows an averaged ROC curve of the 11 individual results using threshold averaging (Fawcett, 2006). The validity of the threshold averaging was ensured by utilizing the same set of thresholds when computing the individual ROC curves.

### 3.2. Single-subject detection

To measure the individual variance of injury regions and assess the potential error introduced by performing group-wise analysis, we employed the 11 rats with TTC staining results. We calculated the FPR and TPR using the infarcted regions from the TTC data and unstable regions from the PET images (as shown in Fig. 2). The results were compared with the FPR and TPR obtained using the infarcted regions and hypo-uptake regions from two-sample *t*-tests. Table 1 presents the overall results.

**Table 1:**
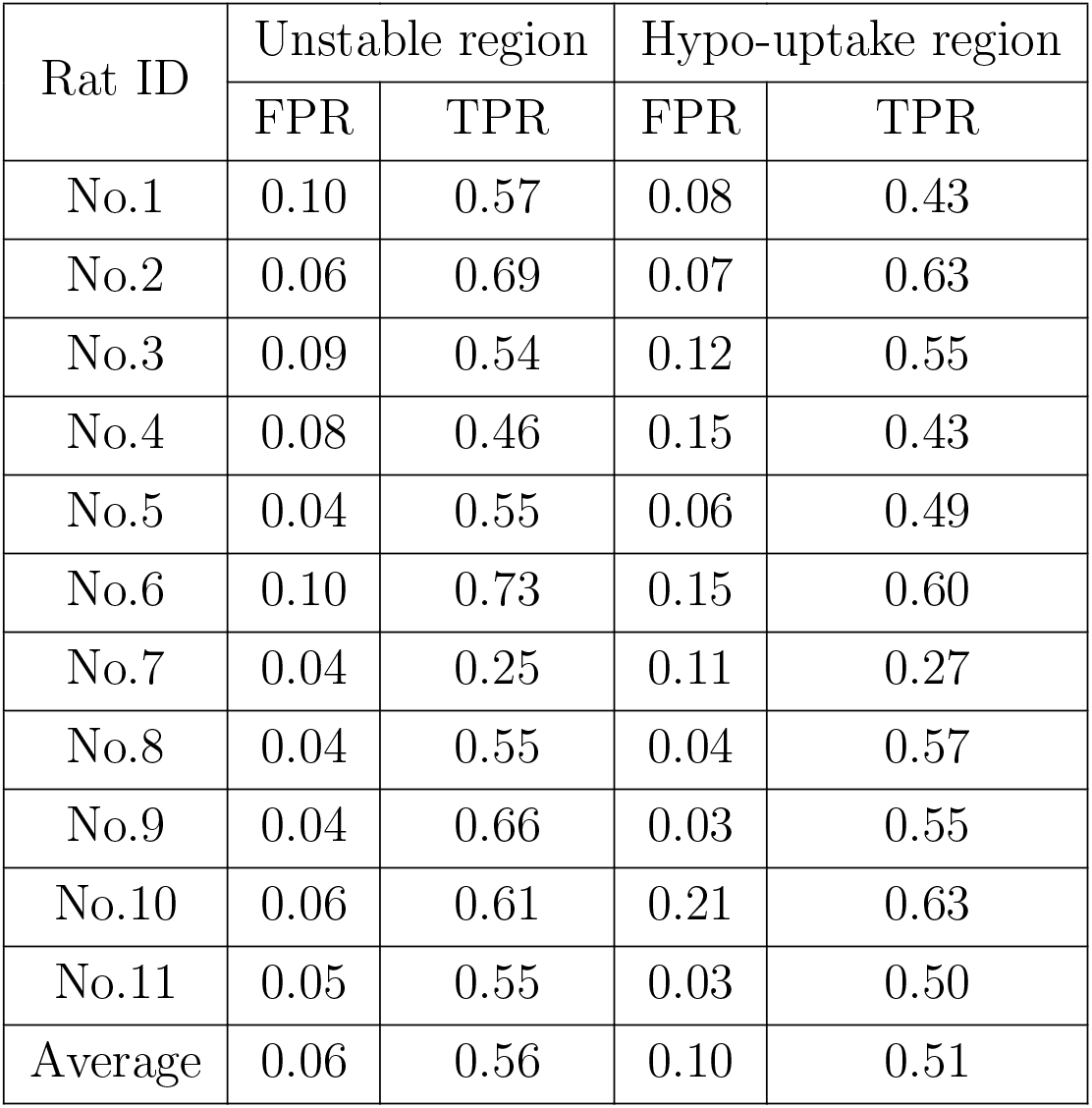
Comparison results between the single-subject analysis (unstable regions using the proposed method) and group-wise analysis (hypo-uptake regions using the two-sample *t*-test with Yakushev’s normalization).

| Rat ID | Unstable region |  | Hypo-uptake region |  |
| --- | --- | --- | --- | --- |
|  | FPR | TPR | FPR | TPR |
| No.1 | 0.10 | 0.57 | 0.08 | 0.43 |
| No.2 | 0.06 | 0.69 | 0.07 | 0.63 |
| No.3 | 0.09 | 0.54 | 0.12 | 0.55 |
| No.4 | 0.08 | 0.46 | 0.15 | 0.43 |
| No.5 | 0.04 | 0.55 | 0.06 | 0.49 |
| No.6 | 0.10 | 0.73 | 0.15 | 0.60 |
| No.7 | 0.04 | 0.25 | 0.11 | 0.27 |
| No.8 | 0.04 | 0.55 | 0.04 | 0.57 |
| No.9 | 0.04 | 0.66 | 0.03 | 0.55 |
| No.10 | 0.06 | 0.61 | 0.21 | 0.63 |
| No.11 | 0.05 | 0.55 | 0.03 | 0.50 |
| Average | 0.06 | 0.56 | 0.10 | 0.51 |

### 3.3. Neurological discoveries

Our proposed method is capable of identifying regions that exhibit high levels of stability or alteration. Regions that display fine linearity between baseline and MCAO images are considered to have minimal influence during ischemia and may be useful as references for studying stroke recovery. In this study, we examined stable regions based on the following criteria.

1. The region should have at least 15 voxels;
2. The median of *R*^2^ is at least 0.9 among all the samples;
3. The 25th percentile is at least 0.8.

The first criterion pertains to the volume size and ensures the robustness of linear regression (Jenkins & Quintana-Ascencio, 2020), as *R*^2^ values can be easily affected by outliers with insufficient data points. The second and third requirements guarantee the prevalence of stable regions among the samples.

Similar to the stable region, an unstable region in the ipsilateral right hemisphere was determined based on the two criteria.

1. The region should have at least 15 voxels;
2. The median of *R*^2^ in the left brain is at least 0.2 greater than that of the same region in the right brain.

Out of the 222 brain regions in the atlas, 194 had at least 15 voxels. We calculated the *R*^2^ value for each region in both hemispheres and obtained their differences (left minus right). The median of the *R*^2^ differences was computed using all subjects from each of the two data sets. A discrimination threshold of 0.2 was chosen for the same reason as described in section 3.1.1. The bar plot in Fig. 5. (a) shows the results of *R*^2^ differences for all 194 regions. The Venn diagrams in Fig. 5. (b) and (c) indicate the number of common stable and unstable regions between the two data sets, respectively.

**Figure 5:**
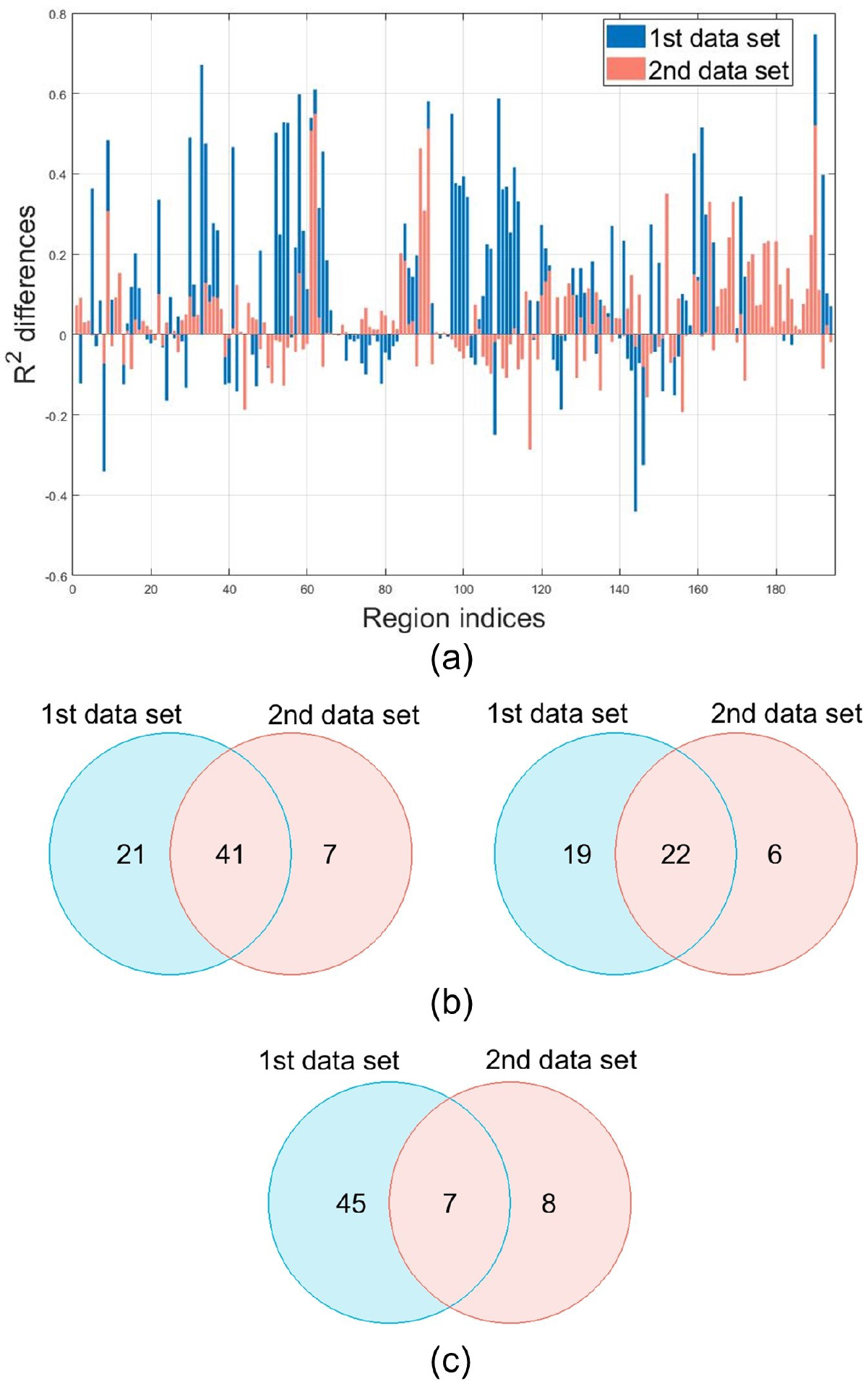
Significant regions can be obtained from the two data sets based on the same criteria. (a) The median of *R*^2^ differences between the left and right brain (left - right) for all subjects in the 194 regions with more than 15 voxels. (b) The Venn diagram showing the number of stable regions in the left brain (left plot) and right brain (right plot) between the two data sets. (c) The Venn diagram showing the number of unstable regions between the two data sets.

Fig. 5. (b) indicates that using the same set of parameters, more stable regions can be calculated from the first data set, including most of the results obtained from the second data set. However, the unstable regions show much less overlap between the two data sets, as shown in Fig. 5. (c). Only seven unstable regions are common between the two data sets, as listed in Table 2. The ventricular system has been observed to undergo positional changes within the ipsilateral hemisphere following MCAO (Lourbopoulos et al., 2012; Walberer et al., 2007). The perirhinal cortex, comprising areas 35 and 36, is a crucial region for sensory processing and cognitive memory. Studies have documented its morphological alterations and abnormalities after stroke. For instance, in a mice stroke model, astrocyte and microglial activation were found to be increased (Schmidt et al., 2015). In another study, the thickness of the perirhinal cortex was significantly reduced in the ipsilesional hemisphere of stroke patients (Haque et al., 2019). Both the primary and secondary auditory areas are constituents of the neocortex and are responsible for processing auditory sensations (Barth et al., 1995; Owjfard et al., 2021). Hearing impairments are frequently observed in stroke patients, and several studies have reported damage in the auditory cortices (Häusler & Levine, 2000; Truong et al., 2012). The temporal association cortex, also located in the neocortex, is situated ventral to the secondary auditory area. Its primary function is recognizing and identifying complex stimuli, and lesions in this region lead to the inability to identify faces (Dahl et al., 2009). Interestingly, regions recognized to have infarction post-ischemia, such as the striatum and hippocampus, were significant in the first data set but failed to meet the threshold in the second data set due to their R2 differences (left - right) lying between the range of [0, 0.2].

**Table 2:** Brain regions having *R*^2^ difference higher than 0.2 from the left to the right brain in both data sets.

| Region indices | Region name | Median of $R^2$ difference | |
| --- | --- | --- | --- |
|  |  | 1st data set | 2nd dataset |
| 9 | Ventricular system | 0.47 | 0.25 |
| 70 | Perirhinal area 35 | 0.50 | 0.47 |
| 71 | Perirhinal area 36 | 0.61 | 0.46 |
| 101 | Primary auditory area | 0.49 | 0.43 |
| 102 | Secondary auditory area, dorsal part | 0.30 | 0.29 |
| 103 | Secondary auditory area, ventral part | 0.61 | 0.51 |
| 218 | Temporal association cortex | 0.75 | 0.48 |

## 4. Discussion

The proposed algorithm in this study was based on the supposition that two primary factors contributed to the disparities observed between scans of the same subject before and after the experimental procedure or disease. The first factor consisted of elements unrelated to the condition, such as the concentration of injected FDG and basal metabolic rate. These factors were not influenced by brain dysfunction or activation; however, they produced intensity ranges that were markedly different. The second factor was due to the experimental conditions, which were typically limited to specific brain regions. Consequently, a regression model was employed to analyze the voxels from the two scans, and the coefficient of determination (*R*^2^) was calculated to ascertain the degree of coherence.

In an ideal scenario where both factors are negligible, and there is no difference between the two scans, it is expected that after spatial normalization, each voxel will correspond to the same anatomical location, and their intensity will match. If a regression model is applied to a specific cluster of voxels, the fitted results will be a straight line with a unit slope. In contrast, when the first factor of differences is present, such as in a sample from the control group, the regression model’s fitted result will also be a straight line, but with a varying slope. The magnitude of the slope indicates how the first factor affects voxel intensities globally. In a more complex scenario where the second factor is also present, the linear relationship between voxel intensities will decrease, and the *R*^2^ value of the regression will decrease from approximately one to a smaller value. Measuring the variation in *R*^2^ can quantify how the voxel intensities deviate from the baseline state, and thus identify significant regions in neurological studies.

The optimal method for intensity normalization in FDG PET quantification has been the subject of a long-standing debate. However, all proposed methods have their advantages and disadvantages. Intensity normalization using the global mean or the contralateral half brain can provide accurate quantification results only when a small part of the brain has metabolic alteration. This is not applicable to stroke analysis as various studies have reported dysfunction and diaschisis from remote regions (Rost et al., 2022; Z. Wang et al., 2020). Reference region normalization has been efficient in studies investigating normal aging (Verger et al., 2021; H. Zhang et al., 2017), Alzheimer’s disease (Nugent et al., 2020), or Parkinson’s disease (Berti et al., 2012; Borghammer et al., 2010). However, a proper reference region in stroke studies is still lacking due to the absence of an atlas of rat brain metabolism after brain ischemia (Nie et al., 2018). The histogram-based normalization method calculates the intensity ratio between each corresponding voxel in an image and an averaged template. The most prevalent ratio is then used as the dividing factor to normalize each image to the template’s intensity range for comparison. This is equivalent to plotting the scatter graph of the image and template. The slope of the line between the origin and each scatter point is then calculated, and the most prevalent slope is selected as the normalization factor. In this sense, the histogram-based method can be considered as a regression approach that fits the voxels individually. However, this voxel-wise pairing can introduce a significant bias when the injury region is profound, and the divided histogram deviates significantly from the ground truth. The limitation of this global method can be resolved by using region-based regression. The new approach utilizing the *R*^2^ value is also more robust when a part of the region has been affected.

To objectively evaluate and compare each quantification method, we utilized ROC in our analyses. Previous studies have shown that hyper-metabolism indicates peri-ischemic regions (or ischemic penumbra), while hypo-metabolic areas represent the ischemic core with tissue fate being infarction on TTC staining (Arnberg et al., 2015; Lou et al., 2007; Yuan et al., 2013). Therefore, ROC analysis was performed on the eleven rats with quantified hypo-uptake regions and their segmented TTC. The advantage of ROC analysis over hypothesis testing is that it is insensitive to the *t*-threshold. The conventional *t*-test used in brain disorders or activation studies requires a carefully determined threshold to detect significant voxels. An underestimated *t*-threshold can generate extra hyper- or hypo-metabolism, while an overestimated *t*-threshold can hide truly significant voxels. However, an ROC curve is generated by a series of FPR and TPR, where each pair is computed from a specific threshold value. Therefore, the final curve indicates the consistency between the target method and the ground truth, and the AUC quantifies the magnitudes. Fig. 4 (b) presents the ROC analysis results based on the eleven rats. For most validation samples, our proposed approach outperforms the four benchmark methods. The second-best method is Yakushev’s cluster-based normalization. It has been suggested that this method requires a careful selection of the significance level, but in ROC analysis, the *t*-threshold is varying, and all methods are comparable. Our results also indicate that for most samples, the GMN, RRN, and HN approaches have similar AUC and performances, which are all inferior to the UR or CN. A mean ROC curve was constructed based on threshold averaging, from which the proposed UR method has the highest sensitivity in most domains.

In addition to its superior sensitivity, our proposed model can identify injury regions in a single-subject manner. Our results have demonstrated both visually (Fig. 2) and quantitatively (Fig. 5) that there is substantial variance in terms of injury location. This variation can be observed among individual samples from the same experiment and across different studies. Therefore, a single-subject model can detect the pathology specifically and provide more accurate results. Table 1 presents the comparison results using the single-subject method and group-wise analysis. In most cases, the individual results of unstable regions have lower FPR and higher TPR than the common results calculated from the 11 rats together.

The proposed method can also identify stable regions believed to have minimal influence under experimental conditions. Among the three criteria used to identify a stable region, the first criterion requires a region to have sufficient voxels to ensure reliable prediction in the regression model. The second criterion imposes a lower limit of *R*^2^ on the average level, while the third criterion ensures the popularity of a large *R*^2^ value. Mean and variance were also tested, but the resulting consistency between datasets was lower than when using the median and percentile. The Venn diagrams in Fig. 5. (b) present the consistency level in both hemispheres. In both sets of data, the left brain has more stable regions than the right brain, which is logically correct since the occlusion was induced in the right brain, and the left part was remote from the focal lesion (Clark et al., 1993; Håberg et al., 2009; Popp et al., 2009). Moreover, despite having fewer stable regions in the second data set, the results are nearly a subset of those from the first data set. This phenomenon suggests that the samples in the second experiment suffer from more severe brain injury, and such batch effect can be observed in relevant MCAO studies (Bunevicius et al., 2013; Schirmer et al., 2023).

Similar to the definition of stable regions, we proposed two criteria to identify an unstable region based on the *R*^2^ difference between the left and right brains (Fig. 5. (a)). Much more unstable regions were identified from the first data set compared to the results from the second data set (Fig. 5. (c)). However, in both sets of data, seven common significant regions were retained, which are listed in Table 2. It was necessary to validate the efficacy of the proposed approach by confirming the biological findings from independent studies. According to our literature search, most of the listed regions were reported to have an anatomical impairment or functional degeneration following brain ischemia.

There are several areas for improvement in future studies. Firstly, it is desirable that the criteria used to identify stable and unstable regions be made adaptable to different datasets. In this study, we adjusted these parameters to select unstable regions based on the TTC infarcted areas. However, our results (as shown in Fig. 5) indicate that using fixed parameters for data from different experiments may lead to non-uniform outcomes. Secondly, MRI scans were not acquired in this study, and hence partial volume correction (PVC) could not be performed. The partial volume effect (PVE) may introduce significant errors to the regression model for small volumetric structures. Therefore, in future works, it would be more accurate to use data with PVC. Thirdly, the proposed method carried out regression on brain regions from a pre-defined atlas. Consequently, the ability to examine more localized areas will be restricted by the deterministic brain region delineation. Further improvements could be accomplished by replacing the region-wise regression with cluster-wise regression using unspecified clusters such as small cubes.

## 5. Conclusion

In this paper, we proposed a regression-based model that can be applied in a single-subject manner to analyze FDG-PET images and study brain metabolism after ischemia. The new method provided accurate identification results, precluding potential errors caused by intensity normalization. The validity of the method was verified by two independent data sets with large sample sizes, and the overall performance was better than the benchmark methods. Our approach was able to detect significant regions in a single-subject manner which can capture and measure inter-subject variations. Apart from MCAO studies, it also has the potential to be applied to other brain disorders via FDG-PET imaging.

## 6. Data and code availability

All the data and code involved in this study are available at: http://yulab.ust.hk/regression/

## 7. Author contributions

Wuxian He: Conceptualization, methodology, software, validation, formal analysis, data curation, writing - original draft, and writing - review and editing.

Hongtu Tang: Investigation, resources, and data curation.

Jia Li: Investigation, resources, data curation, and writing - review and editing.

Xiaoyan Shen: Investigation, resources, and data curation.

Xuechen Zhang: Methodology, software, validation, and visualization.

Chenrui Li: Software, visualization, and writing - review and editing.

Huafeng Liu: Investigation, resources, data curation, supervision, project administration, and funding acquisition.

Weichuan Yu: Conceptualization, methodology, validation, data curation, writing - review and editing, supervision, project administration, and funding acquisition.

## 9. Acknowledgements

This work was supported in part by T12-101/23-N, R4012-18 and C6021-19EF from the Research Grant Council (RGC) of the Hong Kong S.A.R. government of China, MHP/033/20 from the Innovation and Technology Commission (ITC) of the Hong Kong S.A.R., the Hetao Shenzhen-Hong Kong Science and Technology Innovation Cooperation Zone project (HZQB-KCZYB-2020083), the internal fund 3030-009 and BGF.001.2023 from HKUST, the National Natural Science Foundation of China (No: 61525106, U1809204), and the National Key Technology Research and Development Program of China (No: 2017YFE0104000).

## Appendix A. Automatic segmentation and cropping of TTC images

In TTC staining, infarcted areas are distinguished from healthy regions by observing the colors where the infarction is in light yellow while the normal parts are in red. Here we developed an automatic segmentation and cropping tool for picking out the brain slices and possible injury regions to facilitate the research, as shown in Fig.A.6. Firstly, we segmented the brain slices from the noisy background. We found that for TTC images of YCbCr color format, the brain slices always had great intensity differences from the background in the Cr channel. Therefore, by taking both the intensity information and smoothness into account in the Cr channel, we minimized the two-phase Global Chan-Vese (Chan et al., 2006) 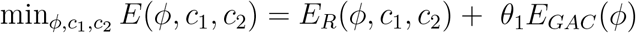 in the level set formulation to segment the brain parts, where the region term *E*_*R*_ and geodesic active contour (GAC) smoothness term (Caselles et al., 1997) read as follows

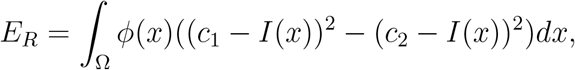

and

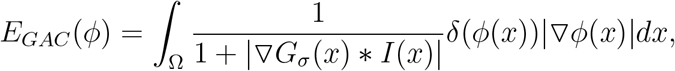

where *ϕ* is the level set function, *c*_1_ and *c*_2_ are constant approximations of the intensity of the background and brain slices respectively, Ω is the image domain, *G*_*σ*_ is the Gaussian smooth kernel of width *σ*, and *θ*_1_ is the weight term.

Secondly, based on the segmentation mask, we used the connected region decomposition algorithm to crop each slice. For the brain slice of each cropped image, we again used the above segmentation functions to segment the infarcted regions in the brain slices.

**Figure A.6:**
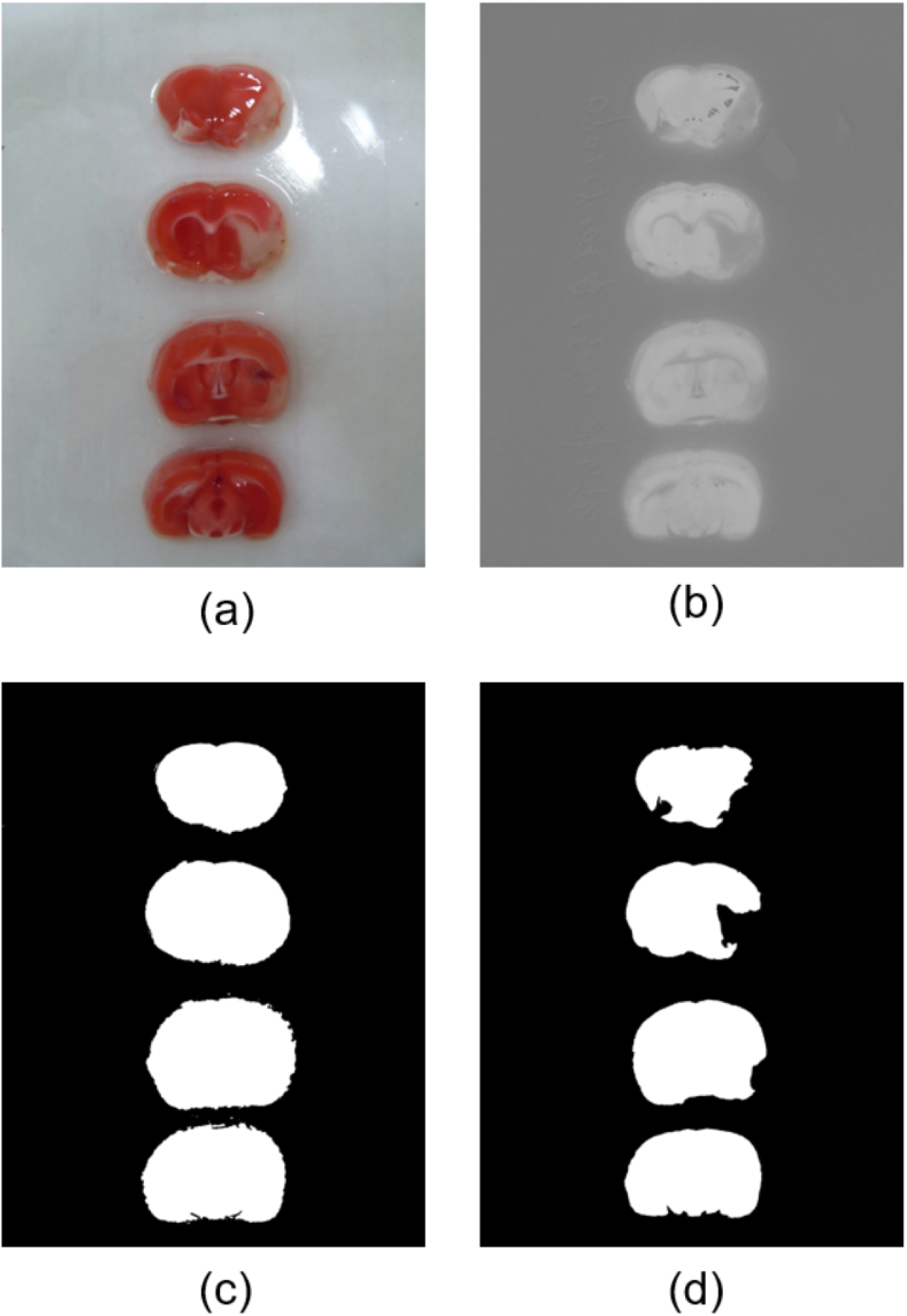
The automatic segmentation pipeline for TTC staining images. (a) An example of an original colored TTC image. (b) The Cr channel of the TTC image, where the noisy background can be easily distinguished from the slices. (c) Level set segmentation results between the slices and background. (d) Level set segmentation of the normal and infarcted regions from the slices. The infarcted regions can be obtained by calculating the difference between (c) and (d).

